# Aquatic insects are dramatically underrepresented in genomic research

**DOI:** 10.1101/2020.08.20.259754

**Authors:** Scott Hotaling, Joanna L. Kelley, Paul B. Frandsen

**Author notes:** Correspondence: Scott Hotaling, School of Biological Sciences, Washington State University, Pullman, WA, 99164, USA, Paul B. Frandsen, Department of Plant and Wildlife Sciences, Brigham Young University, Provo, UT, 84602, USA.

## Abstract

Aquatic insects comprise 10% of all insect diversity, can be found on every continent except Antarctica, and are key components of freshwater ecosystems. Yet aquatic insect genome biology lags dramatically behind that of terrestrial insects. If genomic effort was spread evenly, one aquatic insect genome would be sequenced for every ∼9 terrestrial insect genomes. Instead, ∼24 terrestrial insect genomes have been sequenced for every aquatic insect genome. This discrepancy is even more dramatic if the *quality* of genomic resources is considered; for instance, while no aquatic insect genome has been assembled to the chromosome level, 29 terrestrial insect genomes spanning four orders have. We argue that a lack of aquatic insect genomes is not due to any underlying difficulty (e.g., small body sizes or unusually large genomes) yet it is severely hampering aquatic insect research at both fundamental and applied scales. By expanding the availability of aquatic insect genomes, we will gain key insight into insect diversification and empower future research for a globally important taxonomic group.

**Simple Summary:** Aquatic insects comprise 10% of all insect diversity, can be found on every continent except Antarctica, and are key components of freshwater ecosystems. Yet aquatic insect genome biology lags dramatically behind that of terrestrial insects. If genomic effort was spread evenly, one aquatic insect genome would be sequenced for every ∼9 terrestrial insect genomes. Instead, ∼24 terrestrial insect genomes have been sequenced for every aquatic insect genome. We argue that the limited availability of aquatic insect genomes is not due to practical limitations—e.g., small body sizes or overly complex genomes—but instead reflects a lack of research interest. We call for targeted efforts to expand the availability of aquatic insect genomic resources to gain key molecular insight into insect diversification and empower future research.

## Introduction

There are roughly 1 million described insect species [1]. Of these, ∼100,000 species spend at least one life stage in water [2]. With the rise of high-throughput sequencing, whole genome sequencing has become an increasingly cost-effective research tool [3]. As such, our knowledge of the “genomic natural history” of life has greatly expanded through the combined efforts of individual research groups and large-scale initiatives (e.g., i5K initiative to sequence 5,000 arthropod genomes [4]). Still, while conscious efforts to broadly develop genomic resources across the Tree of Life have been made, major gaps remain. One of these gaps includes the aquatic insects. Despite inhabiting every continent except Antarctica and constituting ∼10% of insect diversity, genomic knowledge of aquatic insects lags far behind terrestrial species. If genomic effort was spread evenly, one aquatic insect genome would be sequenced for every ∼9 terrestrial insect genomes. Instead, ∼24 terrestrial insect genomes have been sequenced for every aquatic insect genome. Here, we show that genomic resources are dramatically limited for aquatic insects relative to terrestrial species in terms of both the number of available genome assemblies and their contiguity, a surrogate for overall quality. We argue that this limitation is not due to any underlying difficulty (e.g., small body size or an unusually large genome) yet it is severely hampering aquatic insect research at both fundamental and applied scales.

With life histories that commonly span aquatic and terrestrial ecosystems, aquatic insects play important ecological roles in many habitats while also providing resource subsidies to higher trophic levels (e.g., mayfly emergence sustaining nesting birds [5]). Aquatic insects are also a global standard for monitoring aquatic ecosystem health [6], a historically organismal approach that is now being enhanced with environmental DNA techniques [7]. The evolution of aquatic insects, however, remains largely a mystery. Depending on the definition used, aquatic insects span at least 12 orders and may include ∼50 separate invasions of freshwater [2]. Five insect orders are almost exclusively aquatic—requiring freshwater for their entire larval development— and include more than 27,000 species: Ephemeroptera (mayflies), Plecoptera (stoneflies), Trichoptera (caddisflies), Odonata (dragonflies and damselflies), and Megaloptera (alderflies, dobsonflies, and fishflies) [8]. The repeated evolution of an aquatic life history raises the question: are insects predisposed to an aquatic lifestyle? But, before this question can be fully addressed, we need a more complete understanding of aquatic insect genome biology.

## Materials and Methods

To test for differences in aquatic and terrestrial genome availability, we used the assembly-descriptors function in the NCBI datasets command line tool to download metadata for all nuclear insect genome assemblies on GenBank (accessed 7 July 2020). Next, we culled the data set to include only the highest quality representative genome for each species based on contiguity and assembly organization (e.g., to the chromosome level). We then determined the life history strategy (aquatic or terrestrial) for each species with a sequenced genome by defining an aquatic insect as any species that spends at least a portion of its larval or adult life stage living and respiring underwater. For our purposes, we chose to exclude the ∼3,500 described species of mosquitoes [9] from our analyses due to their semi-aquatic life cycle where they develop, but do not breathe, underwater [10] and long history in human biomedical research. If we elected to include mosquitoes, they would comprise 61% of all aquatic insect genomes and, a single mosquito genus, *Anopheles*, would account for 51% of the data on its own.

For aquatic and terrestrial insects, we compared the availability and quality of genomic resources in three ways: (1) total number of genomes available, irrespective of contiguity. (2) Number of “highly contiguous” genomes, defined as those with a contig N50 (the mid-point of the contig distribution where 50% of the genome is assembled into contigs of a given length or longer) of 1 Mbp or more following [11]. (3) Number of chromosome-level assemblies (contigs or scaffolds assembled into chromosomes via genetic mapping or similar information) that also exceeded our “highly contiguous” threshold of contig N50 greater than 1 Mbp.

## Results and Discussion

As of July 2020, 536 nuclear insect genomes representing 19 orders have been made publicly available on GenBank (Figure 1; Table S1). Of these, the vast majority are from terrestrial species (*n =* 485), 20 genomes belong to aquatic species, and 31 genomes are from “semi-aquatic” mosquitoes (Figure 1). Aquatic insect genomes comprise just five orders (Diptera, *n* = 5; Ephemeroptera, *n* = 3; Odonata, *n* = 3; Plecoptera, *n* = 3; Trichoptera, *n* = 6), while terrestrial insect genomes span 15 orders (Figure 1).

**Figure 1.**
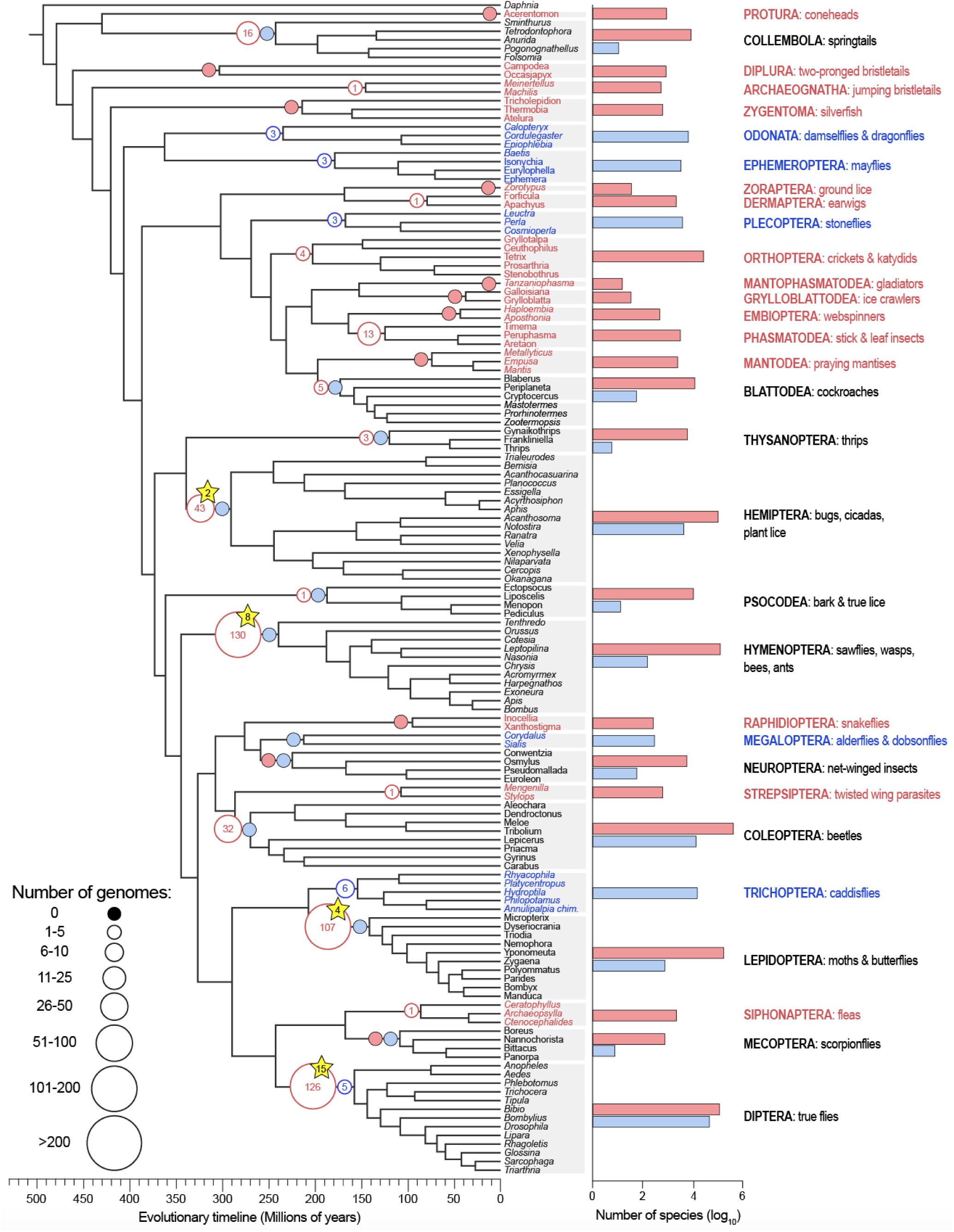
A dated phylogeny of evolutionary relationships among major insect taxonomic groups with the availability of genomic resources for each lineage overlaid. The size of each circle represents the number of available nuclear genomes and their color corresponds to life history strategy, either terrestrial (red) or aquatic (blue). To the right of the tree, the number of described species per group are shown on a log10 scale. Groups that include both terrestrial and aquatic species (e.g., Collembola) are in black font with diversity given for both terrestrial (red) and aquatic (blue) species. Mosquitoes (genomes and species; Order Diptera) were not included in the analysis. Yellow stars indicate the number of chromosome-level assemblies for a given lineage with a contig N50 > 1 million base pairs (there are none for aquatic insects). Species numbers were sourced from a combination of studies [1,2,9,12-16] and the figure was modified from [17]. Complete information of genome availability is provided in Table S1.

Given the total number of insect species that have been described (1,016,507 with mosquitoes excluded) [1] and the number of described aquatic insects (∼100,000) [2], if insect genomes were sampled randomly, nine terrestrial insect genomes would be sequenced for every aquatic insect genome. The reality, however, is that genomic efforts have been dramatically skewed towards terrestrial species (*P*, Fisher’s exact test = 0.0003). To date, 24 unique terrestrial insect genomes have been sequenced for every aquatic insect genome. In other words, if terrestrial insect genome availability was held constant, 33 new aquatic insect genomes (an increase of ∼265%) would need to be made available to bring genomic resources between the groups into balance.

The disparity in genomic resources is even more dramatic when contiguity, our surrogate for total genome *quality*, is considered. Only two aquatic insect genomes (both caddisflies, Order Trichoptera) exceed our “highly contiguous” threshold of a contig N50 > 1 Mbp. This pales in comparison to 56 highly contiguous terrestrial insect genomes spanning five orders (Coleoptera, Diptera, Hemiptera, Hymenoptera, Lepidoptera). More broadly, among the 485 terrestrial insect genomes, the mean contig N50 is nearly 1 Mbp [932.8 thousand base pairs (Kbp)]; for aquatic insects, it’s just 258.5 Kbp. When only highly contiguous (contig N50 > 1 Mbp), chromosome-level assemblies are considered, no aquatic insect genome hits both marks, yet 29 terrestrial insect genomes spanning four orders do (Diptera, Hemiptera, Hymenoptera, Lepidoptera; Figure 1).

Given the substantial contribution of aquatic insects to global insect biodiversity, their importance to ecosystem health and biomonitoring, and the fundamental evolutionary questions they raise, the lack of nuclear genome assemblies for the group is an unfortunate hindrance to research progress in the field. For example, it is impossible to gain a mechanistic understanding of how aquatic insects have repeatedly emerged across the insect Tree of Life until we have properly sampled their genomic diversity.

Some might speculate that while aquatic insects are globally common, they are underrepresented in genomic research because they are small, and therefore difficult to work with, or they have large, unwieldy genomes. To the question of organismal size, given the 16 genomes available for the generally tiny Collembola, including a highly contiguous assembly for *Folsomia candida* [18]— which is just three mm long—organism size is clearly not a limiting factor. And, even if size had historically been limiting, the fact that high-quality reference genomes can now be obtained from single insects (e.g., a mosquito [19]), it certainly is no longer the case. Genome size, however, is less straightforward. For instance, among amphibians, there is a reason that the first frog genome [*Xenopus tropicalis*, 1.7 billion base pairs (Gbp)] [20] was reported ∼8 years before the first salamander genome (*Ambystoma mexicanum*, 32 Gbp) [21]; the latter genome is ∼19x larger and massively more complex. For all insects (including mosquitoes), the mean genome size in the Animal Genome Size Database is 1077 Mbp (*n =* 1,345; accessed 13 July 2020) [22]. While aquatic insects are poorly represented in the Animal Genome Size Database, sequencing-based reports of their genome sizes include five taxonomic orders with a mean size of 600 Mbp (*n =* 20) [23-27]. Thus, there is no evidence that aquatic insect genomes are particularly large and unwieldy when compared to their terrestrial counterparts.

The solution to a lack of aquatic insect genomes is simple: *we should sequence more aquatic insect genomes*. However, to make the best use of resources, we offer two recommendations. (1) Future efforts should first focus on lineages that are relatively speciose for aquatic insects but lack genomic representation. These include alderflies and dobsonflies (Order Megaloptera), aquatic beetles (Order Coleoptera), aquatic true bugs (Order Hemiptera), and aquatic moths (Order Lepidoptera; Figure 1). (2) Since all genome assemblies are not created equal, and contiguity is extremely important for annotating genes and resolving genomic architecture, another focus should be on generating highly contiguous (contig N50 > 1 Mbp), chromosome-level assemblies for aquatic insects, perhaps starting with the five orders that are almost exclusively aquatic (Ephemeroptera, Plecoptera, Trichoptera, Odonata, Megaloptera). By distributing genome sequencing efforts to more properly account for aquatic biodiversity, insect genomics stands to gain considerable insight into the group’s evolution and diversification while simultaneously empowering future research.

## Conclusions

When compared to terrestrial insect genomics, aquatic insects are dramatically underrepresented in genomic research. This underrepresentation is consistent for both the total quantity of available genomes and their quality. This lack of genomic resources is not due to any practical limitation (e.g., body size or genome complexity) and rather appears to simply reflect a lack of interest. We call for targeted efforts to generate more aquatic insect genomes, and particularly for highly contiguous (contig N50 > 1 Mbp), chromosome-level assemblies to be produced. By expanding the availability of aquatic insect genomes, insect and arthropod genome biology stands to gain considerable new potential for research at both fundamental and applied scales.

## Supplementary Materials

Table S1. A table of genome information for all insects used in this study.

## Author Contributions

S.H. and P.B.F. conceived of the study, analyzed the data, and wrote the manuscript. J.L.K. contributed to study design and manuscript preparation. All authors read and approved the final version before submission.

## Acknowledgements

We thank the Kelley and Cornejo Labs at Washington State University and Ellie Armstrong for comments that improved the manuscript.

